# Cellular crowding guides and debundles the microtubule cytoskeleton

**DOI:** 10.1101/830919

**Authors:** A. Z. Płochocka, N. A. Bulgakova, L. Chumakova

## Abstract

Cytoplasm is densely packed with macromolecules causing cellular crowding, which alters interactions inside cells and differs between biological systems. Here we investigate the impact of crowding on microtubule cytoskeleton organization. Using mathematical modelling, we find that only anisotropic crowding affects the mean microtubule direction, but any crowding reduces the number of microtubules that form bundles. We validate these predictions *in vivo* using *Drosophila* follicular epithelium. Since cellular components are transported along microtubules, our results identify cellular crowding as a novel regulator of this transport and cell organization.

---

Distribution of different components inside cells is crucial for cellular, and therefore, organism function. In order for organelles to be delivered to their corresponding biologically relevant locations inside the cell, they are transported via vehicles (motor proteins) along tracks (microtubule cytoskeleton). The microtubules (MTs) forming these tracks are polarized and highly dynamic filaments [1], as their plus-ends undergo dynamic instability. In particular, MTs are either growing or shrinking and can switch between the two states. Despite this highly dynamic behavior of individual MTs, they self-organize into a network, the dynamics of which reaches a steady-state. This steady state is often driven by cell-scale features, e.g. cell geometry and spatial distribution of MT stable minus-ends [2–4].

The properties of the MT network are crucial for cell function. In particular, the mean MT direction is linked to the large-scale direction of transport and cytoplasmic flows [5–7]. The efficacy of intracellular transport additionally depends on the MT bundling, which occurs in many experimental systems [8]. It is defined as the case when two or more MTs are closely apposed, often connected by cross-linking proteins [9]. The presence of bundling promotes the transport by increasing the probability of a motor protein reattachment to a MT upon fall-off [10, 11].

However, the MT network does not exist in isolation, but rather in a crowded cytoplasm densely packed with biopolymers [12]. This dense packing with macromolecules can make the cell interior either isotropic or anisotropic [12–15]. The significance of cytoplasmic crowding is seen in protein folding, where it speeds up transition-limited reactions while slowing down diffusion-limited reactions [13, 16]. Additionally, the crowding creates potential barriers to growing MTs. The only model to date that considers the MTs in the context of crowding analyzes the creation of traffic jams by kinesin-8 [17], whereas the effects of crowding on MTs themselves remain unknown. In this paper we focus on how cellular crowding and its anisotropy affect MT self-organization.

To address this, we combine stochastic simulations, analytical models and *in vivo* experiments. We model cellular crowding as barriers in the cytoplasm, where their positions are either statistically *isotropic* or *anisotropic*, and *homogeneous* or *discrete*. We discover that all barrier types reduce MT bundling, whereas only anisotropic barriers alter their main direction. We validate our predictions *in vivo* using *Drosophila* follicular epithelium at late stages of oogenesis [18–20]. Altogether, we demonstrate that cellular crowding and its directionality impact on the MT network organization and should be considered when studying MT-related processes in cells.

## Model

As cellular crowding is a universal phenomenon, we turn to a system in which MTs can be modelled without excessive oversimplification. In the epithelial tissue, one of the four major tissue types [3], the cortical MTs are restricted to the thin 1*µm* quasi-2d subapical layer (Fig.1a, [2]). This allows to model cells as 2d convex domains, in which MTs grow from points on the boundary *ζ* into the interior (Fig.1b, [21, 22]) at an angle *θ* (or *ϕ*) with respect to the boundary (or the horizontal). All the mathematical model results are presented on elliptical cells, since it is the average cell shape for a given eccentricity [4].

**FIG. 1.**
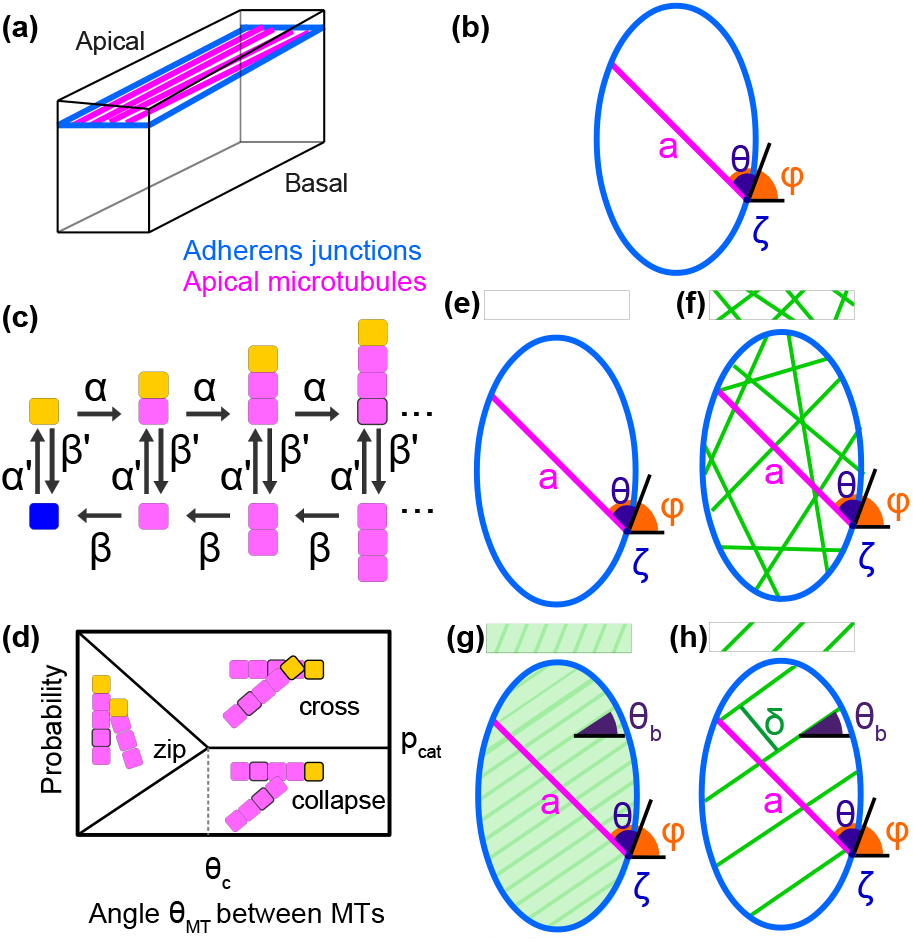
Model setups of the MT dynamics (a-d) and the cytoplasmic crowding (e-h). (a) The apical MTs (*magenta*) in epithelial cells are anchored at the adherens junctions (*blue*) and grow within the 1*µm* layer. (b) A MT growing from the minus-end *ζ* on the boundary (*blue*) into the interior at the angle *θ* (or *ϕ*) with respect to the boundary (or the horizontal); *a* is the cross-section length. (c) Markov chain model of a MT. The rates of polymerization – *α*, catastrophe – *β′*, depolymerization – *β*, and rescue (from either the minus-end (*blue*) or when depolymerizing (*magenta*)) – *α′*. (d) MT interaction: probabilities of a growing MT to collapse, cross, or zip parallel to an existing MT as a function of the angle *θ*_*MT*_ between them. *θ*_*c*_ is the critical angle, *p*_*cat*_ is the probability of catastrophe. (e-h) The four scenarios of crowding barriers (*green*): (e) isotropic homogeneous; (f) isotropic discrete; anisotropic homogeneous cytoplasm with the angle *θ*_*b*_ of anisotropy; and (h) anisotropic discrete barriers at the angle *θ*_*b*_, with spacing *δ*. Boxes indicate labels for the crowding models.

We represent individual MTs as 1d filaments and their dynamic instability via a Markov chain (Fig.1c, [2, 4, 23]), with the of growth *α*, depolymerization *β*, rescue *α′* and catastrophe *β′* (Fig.1c). We set the *base rates* (*α, β, α′, β′*) = (1000, 3500, 4, 1) (as in [4]) and change the catastrophe rate *β′* depending on the nature of barriers. We assume that crowding does not alter the tubulin concentration in the cytoplasm, and hence *α* or *α′*, whereas the depolymerization rate *β* is independent of it [24]. Upon fully depolymerizing, the MT switches to growing at the rescue rate *α′*.

We choose the simplest angle-dependent model of MT interactions (Fig.1d, [2]). When a polymerizing MT encounters an existing one at an angle *θ*_*MT*_, it can grow parallel to it (*zipping*), forming a bundle [25]. Since MTs cannot bend beyond a certain critical angle *θ*_*c*_ due to their rigidity [26], if *θ*_*MT*_ > *θ*_*c*_, the oncoming MT undergoes catastrophe with probability *p*_*cat*_ and crosses otherwise; and for *θ*_*MT*_ < *θ*_*c*_, it collapses, crosses or zips with probabilities 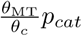, 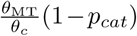, 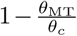 respectively.

To systematically study cellular crowding, we examine four barrier placement scenarios named after the terminology in turbulence. **(1) Isotropic homogeneous** (Fig.1e): the simplified limiting case with small biopolymers, whose distribution is homogeneous and isotropic, is modeled by uniformly increasing the base value of the catastrophe rate *β′*. **(2) Isotropic discrete** (Fig.1f): when the biopolymers are not aligned, but their distribution is not homogeneous, e.g. cortical actin mesh [27], they are modelled as discrete random barriers. Upon encountering a barrier, MTs collapse with the probability *p*_*b*_, increasing the catastrophe rate from *β′* to 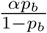. **(3) Anisotropic homogeneous** (Fig.1g): when the biopolymers are aligned, but in the limiting case of being very close to each other, they are modeled as a barrier field at an angle *θ*_*b*_. Here the catastrophe rate 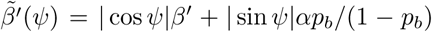 depends on the angle between the MTs and the barriers *ψ* = *ϕ* − *θ*_*b*_, increasing from the base rate *β′* to the 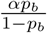 when MTs are perpendicular to the barriers. **(4) Anisotropic discrete** (Fig.1h): The barriers, e.g. actin cables, separated by *δ* are placed at the angle *θ*_*b*_ with respect to the horizontal, and the MTs collapse at barriers with the probability *p*_*b*_. Since the time-scale of the barrier dynamics *in vivo* (e.g. actin cables) is much longer than the MT growth cycle (15sec, [2]), we model them as stationary.

## Microtubule organization

For reported parameter ranges of *β′* ([4] and the references therein), the MT organization is not affected by isotropic crowding (Fig.2a,b), since homogeneous crowding is the limiting case of infinitely close random barriers, and the MT organization is not sensitive to uniformly changing *β′* [4]. Since *β′* has not been measured for crowding scenarios, we investigated increased *p*_*b*_ corresponding to *β′* much higher than the reported range. This progressively weakened the effect of cell geometry [2, 4], reducing MT alignment with the cell major axis (Fig.2a,b *β′* = 5).

**FIG. 2.**
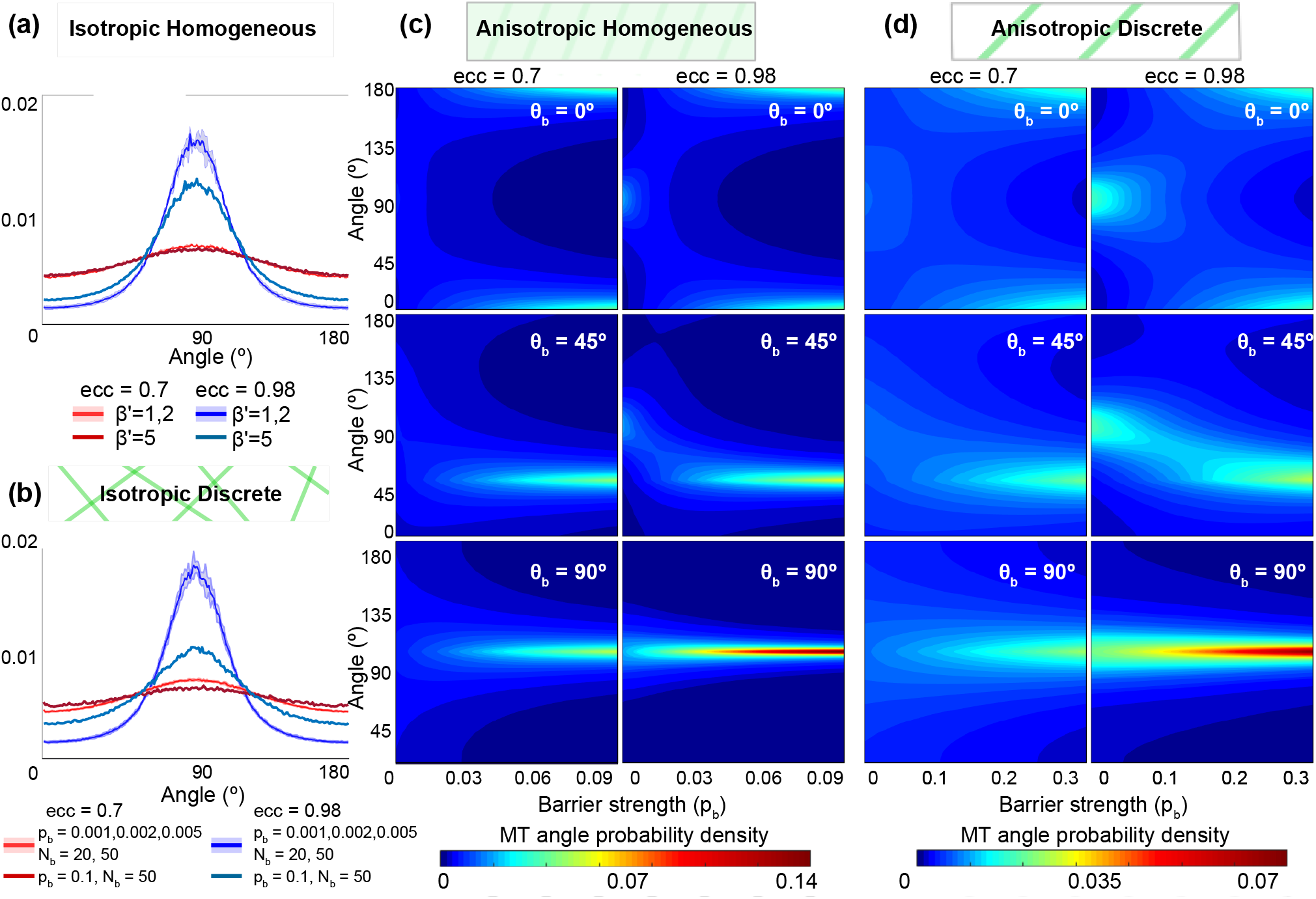
Cellular crowding effect on the MT angle distribution in elongated (*ecc* = 0.98) and non-elongated (*ecc* = 0.7) cells. (a-b) The MT angle distributions for isotropic homogeneous (a) and discrete (b) crowding, for *ecc* = 0.95 (*purple*), and *ecc* = 0.7 (*red*). Robust distributions for the reported values of *β′* = 1, 2, mean (*solid*) and the standard deviation (*envelope*). Reduced effect of cell geometry for *β′* = 5 (*blue curve*). 500 stochastic simulations were run for parameter combination; *p*_*b*_ = 0.001, 0.002, 0.005, 0.1; the number of barriers *N*_*b*_ was varied to keep the barrier density approximately constant: *N*_*b*_ = 20, 50 for *ecc* = 0.7 and *N*_*b*_ = 72, 179 for *ecc* = 0.98. (c-d) Analytic MT angle distributions for anisotropic homogeneous (c) and discrete (D) crowding as a function of the barrier strengths *p*_*b*_ for three barrier angles *θ*_*b*_. In (d) *δ* = 10. The remaining MT instability parameter were kept at their base values.

By contrast, anisotropic crowding introduces competition between the cell geometry and barriers: the former aligns the MTs along the cell major axis, and the latter along the direction of anisotropy (Fig.2c,d). Since the MT angle distribution does not depend on the interaction parameters (*θ*_*c*_, *p*_*cat*_) (see SI, Fig.S1), we used the analytical distribution

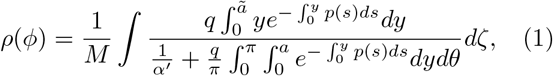

which assumes non-interacting MTs, to analyze its dependence on the barrier strength (for the derivation and the versions for different crowding scenarios see SI section C). Here *M* is the normalization constant, 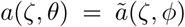 is the cell cross-section, the parameters 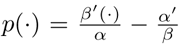 and 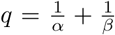, where *β′*(*•*) varies depending on the crowding scenario. For both cases of homogeneous and discrete barriers, we altered the barrier strength *p*_*b*_ for non-elongated and elongated cells (*ecc* = 0.7 and *ecc* = 0.98), while keeping (*α, β, α′*) and (*p*_*cat*_, *θ*_*c*_), constant (Fig.1g,h). For weak barriers, the MT angle distribution is determined by the cell shape, with its peak at the cell major axis angle (90°). With increasing barrier strength, the MTs progressively align with the anisotropy. The rate of this transition depends on the cell geometry and the barrier strength. For elongated cells the effect of the geometry is stronger than for the non-elongated ones, and the MTs align with the cell major axis for larger *p*_*b*_. Since the continuous crowding is the limiting case of infinitely close barriers, the MTs align with anisotropy at smaller *p*_*b*_, comparing to the discrete barrier case (see SI Section D for the study of varying *δ*).

## Validation

We then validated the model predictions *in vivo*. As the strongest effect on MT self-organization is predicted for anisotropic barriers, we used *Drosophila* follicular epithelium, where during late oogenesis (Stage 12, SI Section A) the MTs co-exist with highly aligned densely packed actin cables (Fig.3a,b). In the absence of anisotropic crowding, as in the *Drosophila* embryonic epidermis, MTs orient along the main cell axis [2]. To explore if the actin cables reorient the network, the cells were rotated to have 0° major axis angle. As expected, when not accounted for the actin cable directions, the MT network direction was unbiased (Fig.3c). After flipping the cell images to have the positive angle of actin, the MTs were more likely to have a positive direction (p<0.0001, Fig.3c). This bias was stronger for cells with larger differences between the cell major axis and actin direction (p=0.001 and p=0.0004 for differences above 15° and 25°, Fig.3c). We concluded that actin cables reorient the MT network, and this effect increases with the angle difference between the cell major axis and actin cables.

**FIG. 3.**
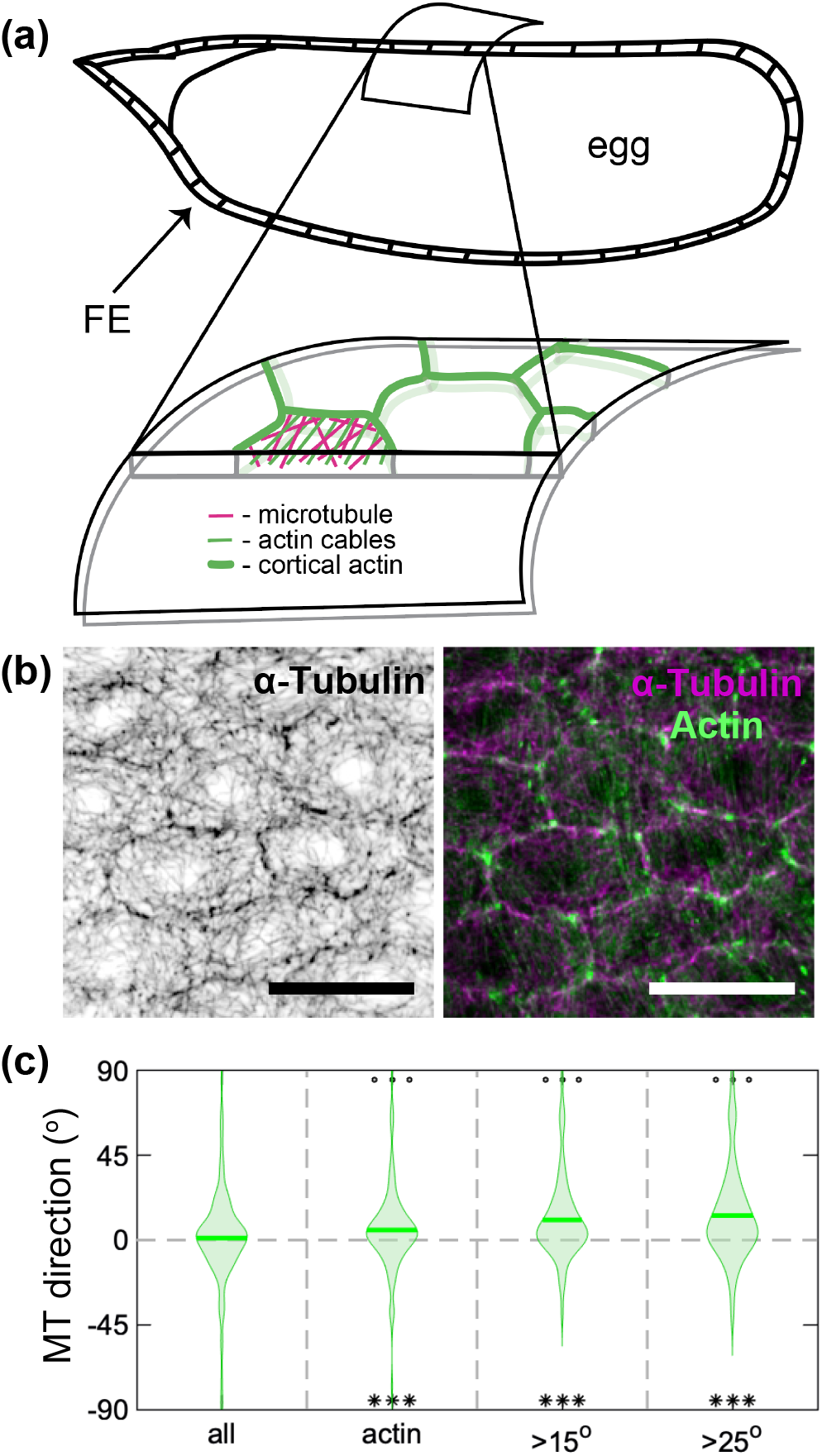
The effect of actin cables in the *Drosophila* follicular epithelium on the MT mean direction. (a) Schematic of the follicular epithelium (FE): a layer of thin cells surrounding the egg chamber with a closer view of FE (bottom): MTs (*magenta*) and actin (*green*). (b) Example of follicular cells stained for MTs (*grey*, left; *magenta*, right) and actin (*green*, right). The scale bar is 10*µ*m. (c) The main direction of MT network without normalization to the direction of actin (all), and with normalization: in all cells (actin), and in cells with the angle between their direction and actin greater than 15°(>15°) and 25° (>25°). ***-p<0.0001 to differ from zero; ^*ooo*^-p<0.001 in comparison to the non-normalized distributions.

## Bundling

To our surprise, upon removal of actin cables by treating ovaries with Latrunculin A the MT organization changed profoundly (Fig.4a, [28]). The MTs appeared more bundled, forming thicker and brighter filaments (Fig.4a), the average area covered by them was reduced (p=0.0005, Fig.4b), while their signal intensity increased (p=0.02, Fig.4c). We concluded that actin cables inhibit bundling *in vivo*.

To explore it further via modelling, we introduced the *bundling factor* as the ratio of MT lengths in bundles to their total length (Fig.4d,e). In all crowding models, the bundling factor was reduced in the presence of barriers (Fig.4f), further decreasing with the overall barrier strength: their number *N*_*b*_ and strength *p*_*b*_ (Fig.4f), and decreased spacing *δ* (SI Fig.S4).

**FIG. 4.**
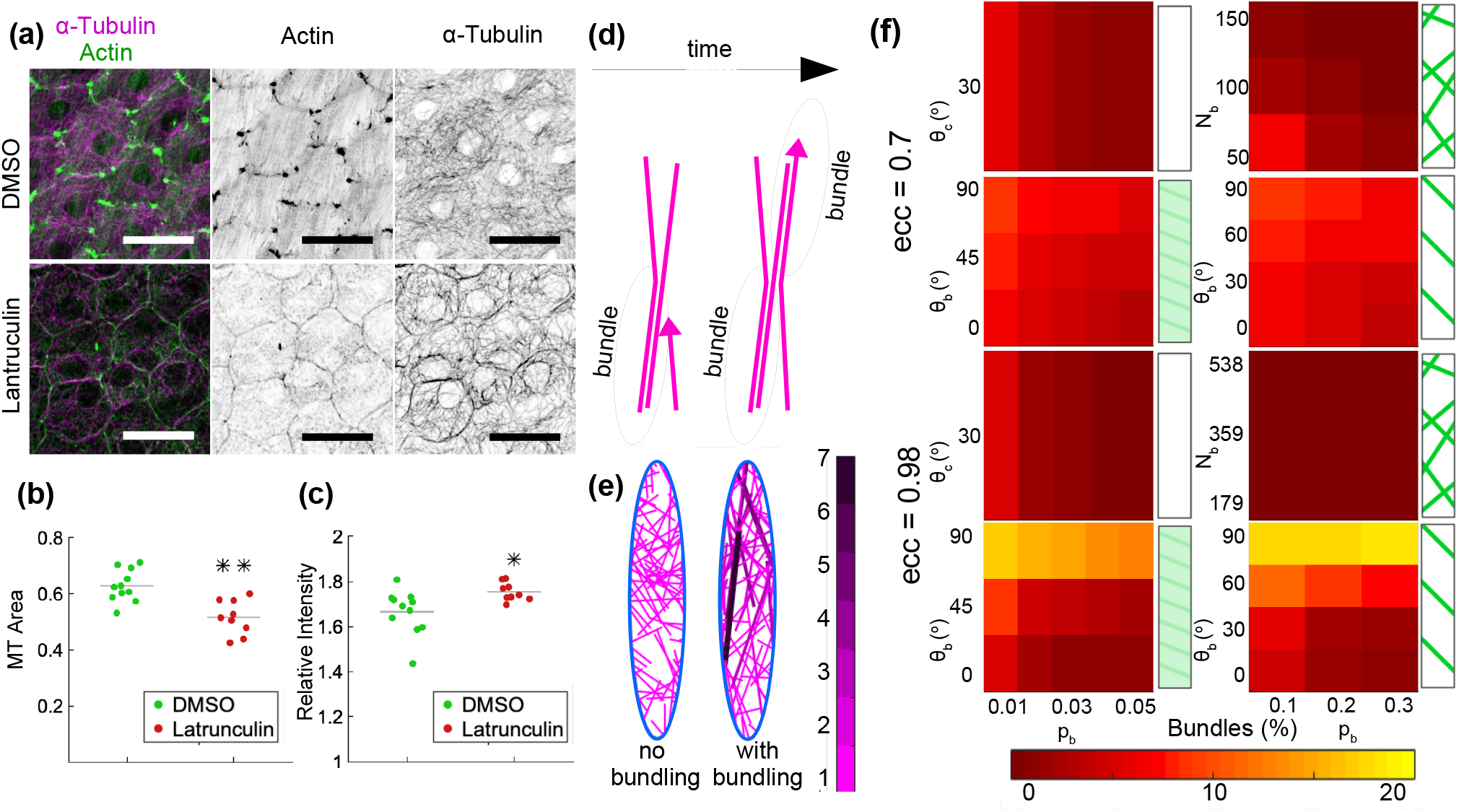
Effect of cellular crowding on MT bundling. (a) *Drosophila* follicular epithelium cells in control (*top*) and with disassembled actin cables after treatment with Latrunculin A (*bottom*), stained for MTs (*magenta* – left, *grey* – right) and actin (*green* – left, *grey* – right). The scale bar is 10*µ*m. (b) Average area covered by MTs (MT signal area divided by the cell area), and (c) signal relative intensity indicating MT bundling. Each dot represents an individual egg chamber in (b) and (c). *-p<0.1, and **-p=0.0005. (d) Bundle formation. (e) Snapshot of stochastic simulations (*ecc* = 0.98, 200 MTs, (*α, β, α′, β′*) = (1000, 3500, 4, 1) with non-bundling (*left*) and bundling MTs (*right*, (*θ*_*c*_, *p*_*cat*_) = (30*°*, 0.01)). (f) Bundling factor (*ecc* = 0.7- *top*, *ecc* = 0.98 – *bottom*), for the four crowding scenarios (clockwise: isotropic homogeneous, isotropic discrete, anisotropic discrete (with *δ* = 10), anisotropic homogeneous) as a function of the barrier strength *p*_*b*_ (*horizontal axis*) and either the number of barriers *N*_*b*_ for the isotropic discrete case, or the angle barrier *θ*_*b*_ for the anisotropic cases (*vertical axis*).

## Conclusion

Here we explored the often overlooked effect of a crowded cytoplasm on MT self-organization. We considered different scenarios using both analytical models and stochastic simulations, and introduced a new measure: MT bundling, by counting MTs which zip along each other. Finally, we validated the model of discrete anisotropic barriers *in vivo* on the *Drosophila* follicular epithelium.

We found that only anisotropic crowding affects the direction of MT network. This is due to the competition between the cell geometry aligning it along the cell major axis [2, 4] and anisotropic crowding redirecting it along itself, where the geometry effect is stronger for more elongated cells. The orientation of the MT network directs intracellular transport [5–7], which in some biological systems is required to be other than the cell major axis. For example, in the follicular epithelium the transmembrane protein Fat2 accumulates along the boundaries parallel to the cell major axis [29]. This localization depends on MTs [19, 29], suggesting the need for their reorientation for the efficient delivery of Fat2 to produce a viable egg. Therefore, cellular crowding anisotropy provides a powerful tool for a cell to redirect the transport and perform its correct function.

We showed both *in vivo* and *in silico* that cellular crowding reduces bundling. How this alters efficacy of intracellular trafficking by molecular motors remains an open question, as bundling can both increase and decrease trafficking by, first, reducing the overall MT density in the cytoplasm, while increasing the probability of motor re-attachment after a fall-of a MT, thus facilitating the cargo reaching the cell boundary. In summary, cellular crowding, though often overlooked, is an important contributor to MT self-organization, and thus to the correct cellular organization and function.

This research was supported by The Maxwell Institute Graduate School in Analysis and its Applications, the Centre for Doctoral Training funded by the UK EP-SRC grant EP/L016508/01, the Scottish Funding Council, Heriot-Watt University and the University of Edinburgh (A.Z.P.); BBSRC BB/P007503/1 (N.A.B.); Royal Society of Edinburgh and the Scottish Government personal fellowship (L.C.); and the Leverhulme trust grant RPG-2017-249 (L.C. and N.A.B).

