## Supplementary information for "Cellular crowding guides and debundles the microtubule cytoskeleton"

**D Sensitivity of the MT self-organization to MT-MT interaction** **6**

**E Competition between the cell geometry and crowding on the mean MT direction** **6**

**F The effect of barrier spacing on the MT organisation for anisotropic discrete crowding** **7**

### A *In vivo* study

#### A.1 Fly stocks, ovary fixation and immunostaining

$w^{1118}$ females were mated and kept at $25^oC$ for 4 days after eclosion. The ovaries were fixed as described in [2] - main text. In brief, the ovaries were dissected in Shields and Sang M3 Insect Medium (S8398, Sigma), incubated for 15 min with 2 $\mu M$ Latrunculin A (Sigma, L5163) or an equivalent volume of DMSO in the M3 Medium, and then immediately fixed in 4% formaldehyde (methanol free, #18814, Polysciences Inc.) in PBS for $20min$ at room temperature. Fixed ovaries were washed three times in PBS with 0.3% Triton X-100, blocked for $1h$ in 5% Native Goat Serum (ab7481, Abcam) in PBS with 0.3% Triton X-100, and incubated with mouse anti-$\alpha$-Tubulin 1:1,000 (T6199, Sigma) primary antibody overnight at $4^oC$. Incubation with the Alexa Fluor 647-conjugated secondary antibodies (1:300, Jackson ImmunoResearch) and Alexa Fluor 488 Phalloidin (1:500, ThermoFisher Scientific) was performed for $2h$ at $25^oC$. Finally, ovaries were dissected into separate ovarioles, muscle sheaths were removed with forceps. The ovarioles were mounted in Vectashield (Vector Laboratories).

##### A.2 Image acquisition and data analysis

Images were acquired using an up-right Olympus FV1000 confocal microscope with a 60x/1.40 NA oil immersion objective. 16-bit images were taken using the FV10-ASW software at a magnification of 0.12 $\mu m/pixel$. For each egg chamber, a 6 z-sections spaced by 0.38 $\mu m$ were imaged, spanning the whole depth of follicular epithelial cells.

Image analysis was done as described in [2] - main text. In short, signal from cortical actin was used to produce cell outlines in the Tissue Analyzer plugin in the Fiji software (https://fiji.sc/). These cell outlines were used to identify each cell as an individual object and fit it to an ellipse to calculate eccentricity and the direction of the ellipse major axis using a script in MATLAB R2018a (Mathworks, www.mathworks.co.uk). The $\alpha$-Tubulin and Phalloidin signals within each cell were filtered using the cell outline as a mask, and the magnitude of the signal according to its direction were calculated to produce histograms of direction gradients [2] - main text. To estimate the degree of microtubule (MT) bundling and signal area of $\alpha$-Tubulin staining, the average projections of the $\alpha$-Tubulin stained stacks were threshold using Otsu's method. The proportion of pixels above threshold relatively to total number of pixels in a cell was used as a measure of signal area. The ratio between mean intensities of pixels above and below threshold was used as a measure of bundling. The script is available at https://github.com/nbul/Cytoskeleton.

##### A.3 Statistical analysis

All *in vivo* data was analyzed using Graphpad Prism 6.0c (www.graphpad.com). Shapiro-Wilk test was used to test the normality of data distributions. For the analysis of effects of actin on the main direction of MT networks, one-way ANOVA with post-hoc Dunn's multiple comparison test was used. The deviation of the mean direction from zero was tested using Wilcoxon Signed Rank Test. Mann-Whitney test was used to compare the areas of MT signal and MT bundling.

#### B Details of the stochastic simulations

We present all the numerical results on elliptical-shape cells, since for cells of a given eccentricity we found the averaged experimental cell shape to be an ellipse ([4] - main text) on elongated ($ecc = 0.98$) and non-elongated ($ecc = 0.7$) cells. The cell minor axis (width) was 60 dimers, and the major axis (height) was adjusted to fit the eccentricity. Each experiment had 200 minus-ends distributed uniformly along the cell boundary.

The parameters of MT dynamics were chosen as follows. The dynamic instability parameters $\alpha, \beta, \alpha', \beta'$ were are within the correspond to experimental parameter ranges (see [4] - main text - for a full discussion on parameter estimation), where the results did not sensitively depend on

| Parameter | Default parameter | Dimensional equivalent |
|---|---|---|
| $\alpha$ | 1000 | $0.15\mu m/s$ |
| $\beta$ | 3500 | $0.52\mu m/s$ |
| $\alpha'$ | 4 | $0.07316s^{-1}$ |
| $\beta'$ | 1 | $0.01829s^{-1}$ |
| $p_{cat}$ | 0.01 | – |
| $\theta_c$ | $30^o$ | – |

Table S1: Summary of key parameters describing MT dynamics in our models.

the dynamical instability parameters. The dimensional equivalents of the parameters are shown in Table S1. The values of MT-MT interaction are not known *in vivo*, and we used $\theta_c = 30^o$ and a low catastrophe of MT upon collision with an existing one ($p_{cat} = 0.01$) so that the MT dynamics simulations closely resembled experimental images in [2] - main text.

Across the four crowding models, the following ranges of parameter values were tested:

**Isotropic homogeneous:** catastrophe $\beta' = 1, 2, 5$ so that the probability of collapse at the barrier is $p_b = 0.001, 0.002, 0.005$;

**Isotropic discrete:** barrier strength $p_b = 0.001, 0.002, 0.005, 0.1$ and $N_b = 20, 50$ for cells of $ecc = 0.7$ and $N_b = 72, 179$ for cells of $ecc = 0.98$;

**Anisotropic homogeneous:** barrier strength $p_b = 0.01, 0.02, 0.03, 0.04, 0.05$, and angle $\theta_b = 0^o, 45^o, 90^o$;

**Anisotropic discrete:** barrier strength $p_b = 0.1, 0.2, 0.3$, angle $\theta_b = 0^o, 30^o, 60^o, 90^o$, and spacing $\delta = 5, 10, 20$.

To collect the statistics of the stochastic simulations, 500 simulations were run for each parameter combination up to the non-dimensional time 10 (equivalent to 550 seconds). The MT angle distribution was averaged over the last 7.5 non-dimensional time-units after the MT network statistics stabilized (see materials and methods in [2] - main text).

### C Analytical MT angle distributions

The analytical MT angle distributions can be derived in the simplified cases when MT are straight non-interacting filaments (Fig.1b-h – main text) growing from a point on the cell boundary $\zeta$ into the cell interior at an angle $\theta$ (or $\phi$) with respect to the cell boundary (or the horizontal). We follow ([4] - main text) by first considering the MT mean survival time. Since *in vivo* and *in silico* the MT angle distribution is length-weighted, we include the possibility of weighting the mean survival time by a function $\gamma(x)$ of the MT length $x$. If $f(x)$ and $g(x)$ are the mean survival time of a MT of length $x$ in the polymerizing and depolymerizing state respectively, then

$$\begin{aligned} f(x) &= (1 - \beta' dt) f(x + \alpha dt) + \beta' dt g(x) + \gamma(x) dt, \\ g(x) &= (1 - \alpha' dt) g(x - \beta dt) + \alpha' dt f(x) + \gamma(x) dt, \end{aligned} \tag{S1}$$

where $dt$ is a small time-increment. Here the terms on the right-hand-side for $f(x)$ (or $g(x)$) are are growing (or shrinking), switching between states, and the weighted time increment. Expanding to leading order in $dt$, we obtain that $f(x)$ and $g(x)$ are governed by

$$\begin{aligned} df(x)/dx &= \beta'/\alpha(f(x) - g(x)) - \gamma(x)/\alpha, \\ dg(x)/dx &= \alpha'/\beta(f(x) - g(x)) + \gamma(x)/\beta, \end{aligned} \tag{S2}$$

and their difference $h(x) = f(x) - g(x)$ satisfies

$$dh(x)/dx = ph(x) - q\gamma(x), \tag{S3}$$

where the two parameters are

$$p(\cdot) = \frac{\beta'(\cdot)}{\alpha} - \frac{\alpha'}{\beta}, \quad \text{and} \quad q = \frac{1}{\alpha} + \frac{1}{\beta}. \tag{S4}$$

For the purposes of this study we allow the catastrophe rate $\beta'(\cdot)$ to depend on the MT length $x$, the position of the minus-end $\zeta$, the growth direction $\theta$ and the growth direction with respect to the barrier angle $\theta_b$, while keeping the other parameters $(\alpha, \beta, \alpha')$ of the dynamic instability constant. We derive the analytic MT angle distribution in the three cases: isotropic homogeneous barriers, anisotropic homogeneous and anisotropic discrete barriers by allowing different functional form of $\beta(\cdot)$.

To find the boundary conditions for Eq.S3, we assume that a MT immediately depolymerizes upon reaching the cell boundary, and hence $g(a) = 0$, $f(a) = h(a)$. Furthermore, since $g(0) = 0$, then $h(0) = f(0)$. The only quantity of interest is the mean survival time, i.e. the mean lifetime of a MT in a growing state starting at length zero, $f(0) = h(0)$. Since it can be both non-length-weighted ($\gamma = 1$) and length-weighted ($\gamma(x) = x$), we denote the corresponding solutions of Eq.S3 as $f_{nw}(0) = h_{nw}(0)$ and $f_w(0) = h_w(0)$:

$$f_{nw}(0) = q \int_0^a e^{-\int_0^y p(s)ds} dy, \quad f_w(0) = q \int_0^a y e^{-\int_0^y p(s)ds} dy. \tag{S5}$$

At the cell level, these lifetimes are also a function of the MT minus-end position $\zeta$ and angle $\theta$. Finally, the MT angle distribution in the full cell is the ratio of the length-weighted to the non-length weighed full lifetimes, integrated over the cell boundary

$$\rho(\phi) = \frac{1}{M} \int \frac{q \int_0^{\tilde{a}} y e^{-\int_0^y p(s)ds} dy}{\frac{1}{\alpha'} + \frac{q}{\pi} \int_0^{\pi} \int_0^a e^{-\int_0^y p(s)ds} dy d\theta} d\zeta, \tag{S6}$$

where $M$ is the normalization constant, the cell cross-section length is denoted as $a(\zeta, \theta)$ and $\tilde{a}(\zeta, \phi)$ depending on the arguments, and the MT minus-end density is taken to be uniform on the cell boundary.

We now consider the three cases of the barrier distributions, isotropic homogeneous, anisotropic homogeneous and anisotropic discrete, and how it affects the catastrophe rate $\beta'(\cdot)$.

#### C.1 Isotropic homogeneous barriers

In the case of the homogeneous isotropic barriers, $\beta' = const$, and hence the MT angle distribution Eq.S6 is

$$\rho(\phi) = \frac{1}{M} \int \frac{1 - e^{-\tilde{a}p}(1 + \tilde{a}p)}{1 + \frac{q\alpha'}{\pi} \int_0^{\pi} \frac{1 - e^{-ap}}{p} d\theta} d\zeta, \tag{S7}$$

as in [4] - main text, where $p = \frac{\beta'}{\alpha} - \frac{\alpha'}{\beta}$ and $q = \frac{1}{\alpha} + \frac{1}{\beta}$ are constant.

Since *in vivo* typically the rates of growth and polymerization are much higher than those of catastrophe and rescue, $\alpha, \beta \gg \alpha', \beta'$, the MT angle distribution is not sensitive to moderate changes in $\beta'$ and asymptotically $\rho(\phi) \sim \int a^2 d\zeta$ ([4] - main text).

#### C.2 Anisotropic homogeneous barriers

We model the anisotropic homogeneous crowding as a field of barriers with the barrier strength $p_b$ at an angle $\theta_b$ with respect to the horizontal (Fig.1g – main text). In the direction of the barrier field, the catastrophe rate remains at its base value $\beta'$. The catastrophe rate increases with the angle $\psi = \phi - \theta_b$ between the MT ($\phi$) and the barrier field ($\theta_b$), reaching the maximum $\frac{\alpha p_b}{1 - p_b}$ in the direction normal to the barrier field, ensuring that the probability of catastrophe at the barrier is

$p_b$. The simplest parameterization of the catastrophe rate as a function $\psi$ is

$$\tilde{\beta}'(\psi) = |\cos\psi|\beta' + |\sin\psi|\left(\frac{\alpha p_b}{1-p_b}\right), \tag{S8}$$

where $\beta'$ is the base rate. Hence the parameter $p(\cdot) = p(\psi)$ in Eq.S4 becomes

$$p(\psi) = \frac{\tilde{\beta}'(\psi)}{\alpha} - \frac{\alpha'}{\beta}, \quad \text{where} \quad \psi = \phi - \theta_b, \tag{S9}$$

leading to the MT angle distribution

$$\rho(\phi) = \frac{1}{M}\int \frac{p^{-2}(1 - e^{-\tilde{a}p}(1+\tilde{a}p))}{1 + \frac{q\alpha'}{\pi}\int_0^\pi \frac{1-e^{-ap}}{p}d\theta}d\zeta, \tag{S10}$$

where $M$ is the normalization constant, and the cell cross-section is denoted as $a(\zeta,\theta)$ or $\tilde{a}(\zeta,\phi)$ depending on its arguments.

#### C.3 Anisotropic discrete barriers

To model anisotropic discrete crowding, we place parallel barriers at an angle $\theta_b$ with the spacing $\delta$ between the barriers. When a MT encounters one of the anisotropic discrete barriers, the catastrophe rate $\beta'$ increases to $\frac{\alpha p_b}{1-p_b}$, so that the probability of collapse at the barrier is $p_b$. For individual MT growing from the minus-end $\zeta$ at the angle $\theta$ with respect to the cell boundary (Fig.1h – main text), the perceived spacing between the barriers is $L(\zeta,\theta)$, and since the barriers are relatively thin we use a simplifying assumption that their perceived width is $2\varepsilon \ll \delta$.

The increase of the parameter $p$ at the barriers is modeled as jumps from the base value $p_0 = \frac{\beta'}{\alpha} - \frac{\alpha'}{\beta}$ to $p = p_0 + p_1$ that ensures the correct increase in the catastrophe rate $\beta'$

$$p = p_0 + p_1 \sum_{k=1}^{N(x,\zeta,\theta)} \left[H(x - kL(\zeta,\theta) + \varepsilon) - H(x - kL(\zeta,\theta) - \varepsilon)\right]. \tag{S11}$$

Here $p_1 = \left(\frac{\alpha p_b}{1-p_b} - \beta'\right)\frac{\beta'}{\alpha}$, $N(x,\zeta,\theta)$ is the number of barriers the MT of length $x$ growing from $\zeta$ at an angle $\theta$ has passed, and $H(\cdot)$ is the Heaviside function. Therefore,

$$\int_0^y p(s)ds = p_0 y + p_2 N(y,\zeta,\theta), \tag{S12}$$

where $p_2 = 2\epsilon p_1 = 2\epsilon\left(\frac{\alpha p_b}{1-p_b} - \beta'\right)\frac{\beta'}{\alpha}$. The non-length-weighted and length-weighted MT survival times are therefore

$$f_{nw}(0) = q\int_0^a e^{-p_0 y - p_2 N(y,\zeta,\theta)}dy, \tag{S13}$$

$$f_w(0) = q\int_0^a e^{-p_0 z - p_2 N(z,\zeta,\theta)}z dz. \tag{S14}$$

Hence, the MT angle distribution $\rho(\phi)$ is

$$\rho(\phi) = \frac{1}{M}\int \frac{\int_0^{\tilde{a}} e^{-p_0 z - p_2 \tilde{N}} z dz}{1 + \frac{q\alpha'}{\pi}\int_0^{\pi}\int_0^{a} e^{-p_0 z - p_2 N} dz d\theta} d\zeta, \tag{S15}$$

where $M$ is the normalization constant, depending on the arguments, the cell cross-section $a$ is denoted as $a(\zeta,\theta)$ or $\tilde{a}(\zeta,\phi)$, and the number of barriers $N$ that a MT has encountered as either $N(z,\zeta,\theta)$ or $\tilde{N}(z,\zeta,\phi)$, depending on its arguments.

### D Sensitivity of the MT self-organization to MT-MT interaction

We found that the MT angle distribution is not sensitive to the MT-MT interaction, parameterized by $(\theta_c, p_{cat})$, for the anisotropic barrier cases (both homogeneous and discrete) (Fig.S1), since the MT angle distribution in the full stochastic simulations with interacting MTs agree with the analytic distributions in Eq.S10 and Eq.S15, derived for non-interacting MTs.

### E Competition between the cell geometry and crowding on the mean MT direction

Here we present additional studies of the ability of cellular crowding to reorient MT network. We confirm in Fig.S2 that the isotropic crowding scenarios (both homogeneous and discrete) do not alter the mean of the MT angle distribution (Fig.S2, top row) regardless of the cell elongation. For anisotropic crowding scenarios (Fig.S2, bottom row), the mean of the MT angle distribution progressively shifts to the angle of anisotropy $\theta_b$ with the increasing overall barrier strength (their

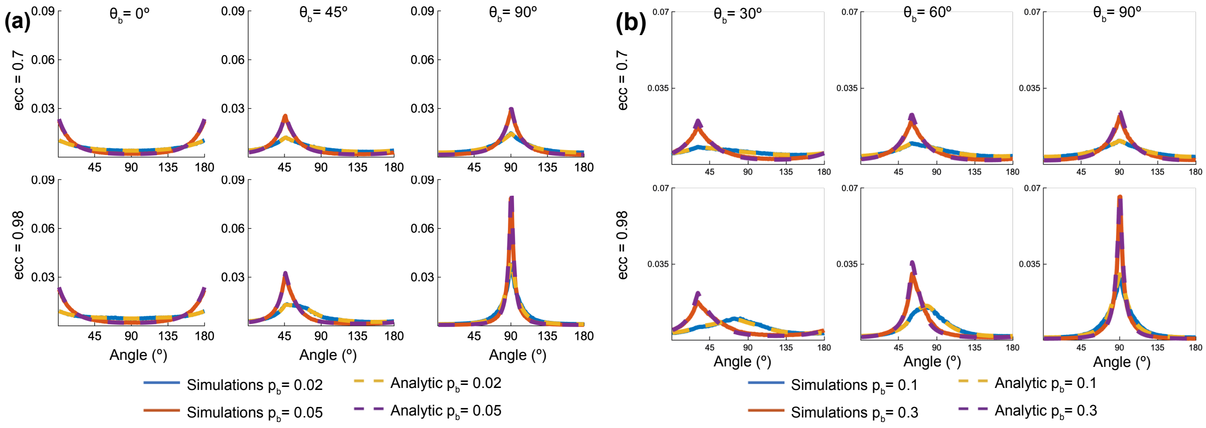

Figure S1: MT angle distribution is not sensitive to the MT-MT interaction parameters. (a) Anisotropic homogeneous barriers: the MT angle distribution in the stochastic simulations with interacting MTs and the analytical MT angle distribution (Eq.S10) for non-interacting MTs, for different barrier angle $\theta_b = 0^o, 45^o, 90^o$, in both elongated and non-elongated cells ($ecc = 0.7$ and 0.95). (b) Anisotropic discrete barriers: the MT angle distribution in the stochastic simulations with interacting MTs and the analytical MT angle distribution (Eq.S15) for non-interacting MTs, for different barrier angle $\theta_b = 0^o, 45^o, 90^o$, in both elongated and non-elongated cells ($ecc = 0.7$ and 0.95). For both (a,b), the number of MTs 200, number of simulations 500 for each parameter set, $(\alpha, \beta, \alpha')$ at base values $(1000, 3500, 4)$, $(\theta_c, p_{cat}) = (30^0, 0.01)$.

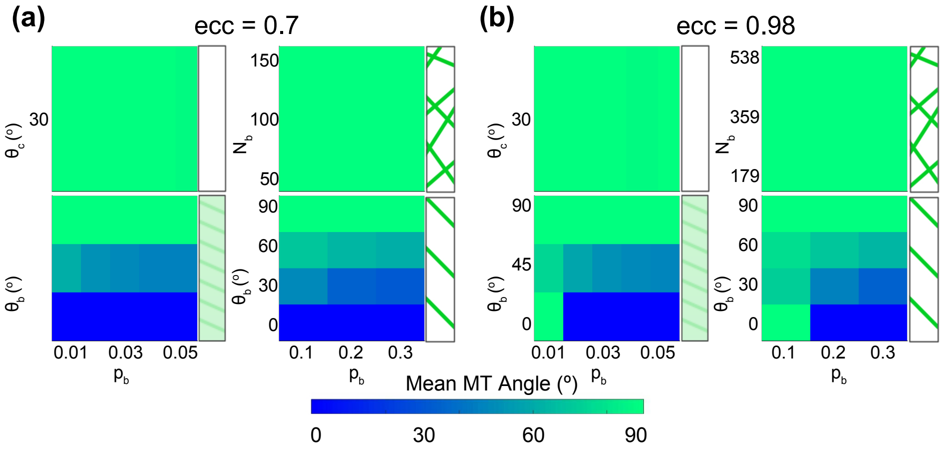

Figure S2: The mean of the MT angle distribution as a function of the barrier strength ($p_b$, $N_b$) and direction $\theta_b$ for two cell elongations, (a) $ecc = 0.7$ and (b) 0.98, and the four crowding scenarios (clockwise: isotropic homogeneous, isotropic discrete, anisotropic discrete, anisotropic homogeneous). All the MT interaction and dynamical instability parameters (except $\beta'$) were kept at their base values.

strength $p_b$ and number $N_b$). However, for the same barrier parameters, the effect of the elongated cells ($ecc = 0.98$ as compared to $ecc = 0.7$) to orient the MTs along the cell major axis ($90^o$) persists until a higher barrier strength $p_b$. This again confirms that in the competition between the cell geometry and anisotropic crowding to reorient MT network, the effect of cell geometry is stronger in more elongated cells.

### F The effect of barrier spacing on the MT organisation for anisotropic discrete crowding

We additionally consider how the barrier spacing affects the directionality of MTs.

First, consider the full MT angle distributions in Fig.S3. For low barrier strength the MTs are aligned with the cell major axis ($90^o$), and as the barrier strength $p_b$ increases, the MTs become progressively more aligned with the barrier angle $\theta_b$. However, since the denser barriers have a stronger inhibiting effect on the MT dynamics than the less dense ones ($\delta = 5$ in Fig.S3a as compared to $\delta = 20$ in Fig.S3b), they reorient MTs at lower barrier strengths $p_b$. We conclude that in cells with denser actin cables, the MTs would be more likely to follow the actin cables direction whereas in cells with less dense cables, the MTs would align more in accordance with the cell geometry.

Finally, we consider the effect of discrete barriers (e.g. actin cables) on the MT mean direction and bundling Fig.S4. Fig.S4a supports main findings that weaker barriers (here – the smaller barrier density), lead to the stronger effect of cell geometry guiding the mean MT direction to be along the cell major axis. As for the bundling factor (Fig.S4b), this additional study supports out main findings that any crowding inhibits bundling. That said, the bundling is higher when the MT dynamics is less interrupted – either when the barriers are parallel to the main cell axis, or the barrier strength is small (smaller $p_b$ or lager spacing $\delta = 20$ between the barriers). Additionally, when bundles form, longer bundles persist for longer periods of time compared to the short bundles. Since elongated cells allow for longer bundles, as compared to less elongated cells, the bundling factor

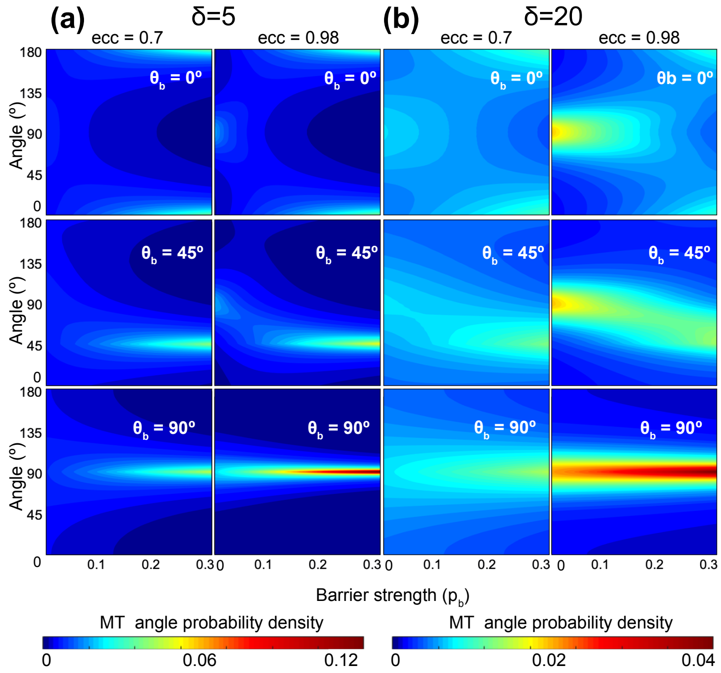

Figure S3: MT angle distributions (*vertical*) for anisotropic discrete barriers for two different barrier spacing (a) $\delta = 5$ and (b) $\delta = 20$, as the function of the barrier strength $p_b$. The MT interaction and dynamic instability parameters (except for $\beta'$) were kept at their base values.

186 for them is higher, making MT bundles in them more resistant to the inhibiting effects of crowding.

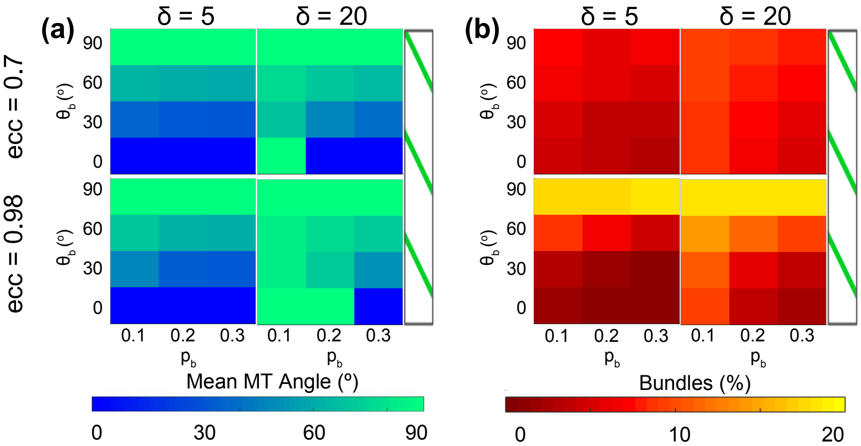

Figure S4: The effect of the barrier spacing on MT mean direction and bundling for anisotropic discrete barriers. The mean MT direction (a) and the bundling factor (b) are shown as a function of the barrier strength $p_b$ and the anisotropy angle $\theta_b$ for two barrier spacings ($\delta = 5$, *left*, and $\delta = 20$, *right*) for less elongated ($ecc = 0.7$, *top*) and more elongated ($ecc = 0,98$, *bottom*) cells.
